# Time-restricted feeding ameliorates MCDD-induced steatohepatitis in mice

**DOI:** 10.1101/2023.11.15.567214

**Authors:** Ik-Rak Jung, Rexford S. Ahima, Sangwon F. Kim

**Author notes:** Corresponding Authors Sangwon F. Kim Rexford S. Ahima.

## Abstract

Non-Alcoholic Steatohepatitis (NASH) is an inflammatory form of Non-Alcoholic Fatty Liver Disease (NAFLD), closely associated with disease progression, cirrhosis, liver failure, and hepatocellular carcinoma. Time-restricted feeding (TRF) has been shown to decrease body weight and adiposity and improve metabolic outcomes, however, the effect of TRF on NASH has not yet been fully understood. We had previously reported that inositol polyphosphate multikinase (IPMK) mediates hepatic insulin signaling. Importantly, we have found that TRF increases hepatic IPMK levels. Therefore, we investigated whether there is a causal link between TRF and IPMK in a mouse model of NASH, i.e., methionine and choline deficient diet (MCDD)-induced steatohepatitis. Here, we show that TRF alleviated markers of NASH, i.e., reduced hepatic steatosis, liver triglycerides (TG), serum alanine transaminase (ALT) and aspartate aminotransferase (AST), inflammation and fibrosis in MCDD mice. Interestingly, MCDD led to a significant reduction in IPMK levels, and the deletion of hepatic IPMK exacerbates the NASH phenotype induced by MCDD, accompanied by increased gene expression of pro-inflammatory chemokines. Conversely, TRF restored IPMK levels and significantly reduced gene expression of proinflammatory cytokines and chemokines. Our results demonstrate that TRF attenuates MCDD-induced NASH via IPMK-mediated changes in hepatic steatosis and inflammation.

## Introduction

Nonalcoholic steatohepatitis (NASH) is a severe progression of nonalcoholic fatty liver disease (NAFLD), characterized by hepatic Triglyceride (TG) accumulation, inflammation, liver fibrosis, and hepatocellular injury, which can progress to cirrhosis and hepatocellular carcinoma (HCC) (Loomba et al., 2021). Numerous studies have proposed mediators that regulate NASH initiation and progression, including insulin resistance (Chitturi et al., 2002), oxidative stress (Masarone et al., 2018), mitochondrial dysfunction (Begriche et al., 2006), cell death (Kanda et al., 2018), immune cell activation (Huby & Gautier, 2022), and genetic/epigenetic factors (Jonas & Schürmann, 2021). Lifestyle modifications, such as reducing calorie intake, exercise, and weight loss are recommended for the treatment of NAFLD/NASH (Baumeier et al., 2015; de Castro-de-Paiva et al., 2022; Golbidi et al., 2012; van der Windt et al., 2018). However, there are few effective therapies to address the increasing global prevalence of NAFLD/NASH (Oseini & Sanyal, 2017). Time-restricted feeding (TRF) is a dietary intervention that involves consuming food within a consistent daily time window, typically spanning 8 to 10 hours, without the need to reduce overall calorie intake (Rynders et al., 2019). Several studies have reported favorable outcomes associated with TRF, including reductions in body weight and adiposity, improvements in blood pressure, and enhanced insulin sensitivity in both human and rodent studies (Chaix et al., 2014; Hatori et al., 2012; Sutton et al., 2018). While the precise mechanisms remain unexplored, it has been shown that TRF prevents the development of liver steatosis and NASH when fed a high-fat diet (Hatori et al., 2012) or a high-fat high-sucrose diet (Shu et al., 2022). Inositol polyphosphate multikinase (IPMK) is responsible for producing inositol tetrakisphosphate (IP4) and inositol pentakisphosphate (IP5) (Frederick et al., 2005). Additionally, IPMK exhibits PI3K activity (Maag et al., 2011) and plays a pivotal role in various cellular signaling pathways. IPMK functions as an adaptor protein, interacting with key proteins, including a mechanistic target of rapamycin (mTOR), AMP-activated protein kinase (AMPK), serum response factor (SRF), and Unc-51-like autophagy-activating kinase (ULK) (Bang et al., 2012; Guha et al., 2019; E. Kim et al., 2013; S. Kim et al., 2011). Recent research has uncovered IPMK’s significant role as a key regulator of Toll-like receptor (TLR) signaling, particularly in myeloid cells, especially macrophages (E. Kim et al., 2017; H. Lee et al., 2023). In this context, IPMK modulates the production of inflammatory cytokines by regulating TRAF6 protein stability through direct interaction with TRAF6 and preventing K48-linked ubiquitination. IPMK has been shown to play a critical role in liver inflammation (Guha et al., 2019). We have reported that IPMK enhances hepatic insulin sensitivity and suppresses gluconeogenesis (Jung et al., 2022). In preliminary studies, we found that TRF increased IPMK expression in liver.

In this study, we show that time-restricted feeding (TRF) improves steatosis and inflammation in a mouse model of NASH, i.e., mice fed a methionine-choline deficient diet (MCDD). We found that IPMK protein was decreased in the liver of MCDD mice. To determine whether the loss of IPMK exacerbates the NASH phenotype, we fed wild-type (WT) and liver-specific IPMK knockout (LKO) mice a MCDD, and assessed liver histology, hepatocellular injury, and inflammation. Hepatic IPMK deficiency worsened hepatic steatosis, elevated serum ALT and AST levels and expression of inflammatory genes in mice fed MCDD, suggesting a role as a molecular mediator of the effect of TRF in NASH.

## Results

### IPMK is decreased in the liver of NASH mice

We previously reported that IPMK protein level was decreased in mice fed a high fat diet (HFD) (Jung et al., 2022). To determine whether hepatic IPMK expression is affected during NASH progression, we measured IPMK protein levels in the liver of C57BL/6J mice fed MCDD. Similar to findings from other studies (Machado et al., 2015; Rizki et al., 2006), the MCDD significantly reduced body weight (Fig. 1A) and increased hepatic TG, serum ALT and AST (Fig. 1B-D). These changes were consistent with the observed increase in hepatic steatosis and inflammation as shown in the histological analysis (Fig. 1E). Fibrosis was also increased in MCDD mice (Fig. 1E). The expression levels of pro-inflammatory cytokines, such as TNFα, significantly increased, along with a substantial upregulation of pro-inflammatory chemokines (MCP1, CCL3, CCL4, and CCL5), and fibrosis related genes (Col1a1, TIMP1 and TGFβ) in the liver of MCDD mice (Fig. 2A-L). Notably, both IPMK protein levels (Fig. 1F) and mRNA expression (Fig. 2A) were significantly lower in the liver of MCDD mice compared to mice on control diet (ND-fed mice).

**Figure 1.**
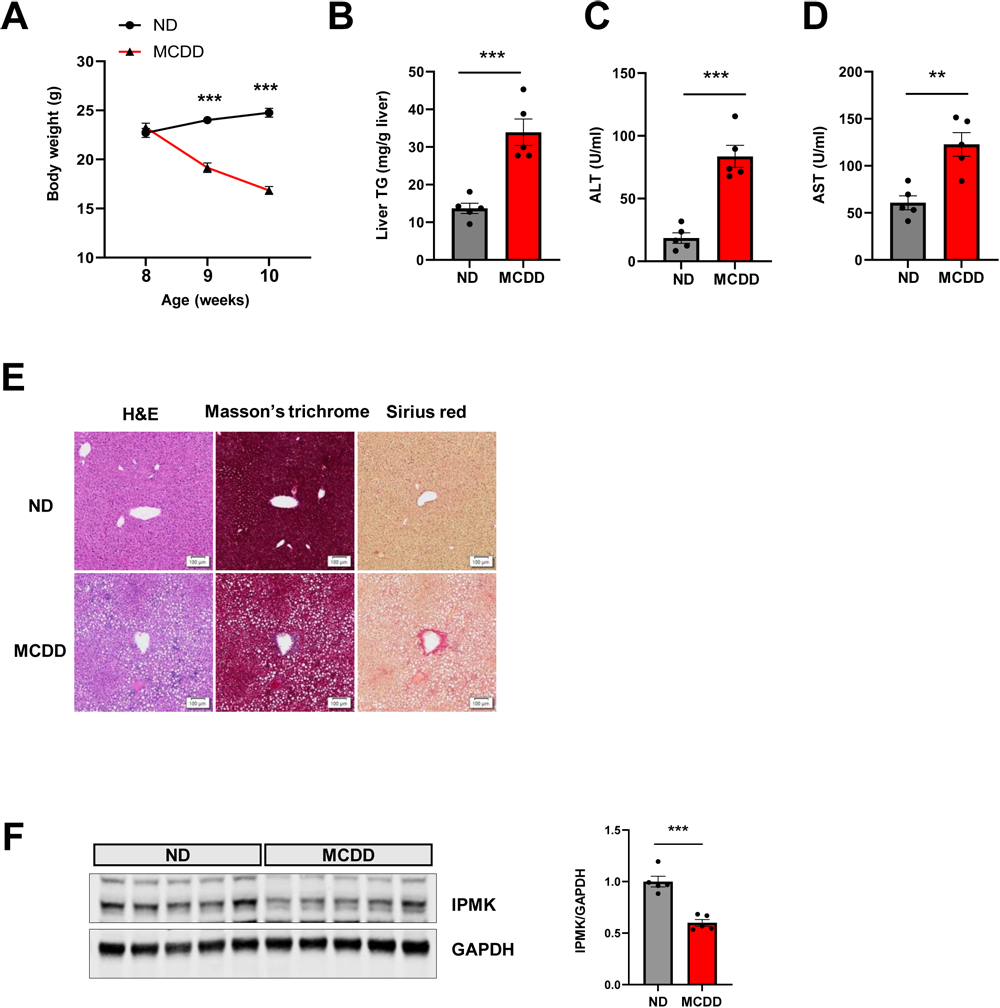
IPMK protein is decreased in the liver of MCDD mice. (A) Body weight of ND (n = 5) and MCDD (n = 5) fed mice was measured weekly from age 8 to 10 weeks. (B) Liver TG, serum (C) ALT and (D) AST were measured to evaluate the levels of liver injury. (E) Liver paraffin sections were stained with H&E, Masson’s trichrome, and Sirius red to determine the levels of steatosis, inflammation, and fibrosis (scale bar, 100 um). (F) Protein levels of IPMK in the liver from ND or MCDD-fed mice were analyzed by immunoblotting (n = 5 per group). Levels of IPMK protein were quantified. Data are mean ± SEM; **p < 0.01, ***p < 0.001.

**Figure 2.**
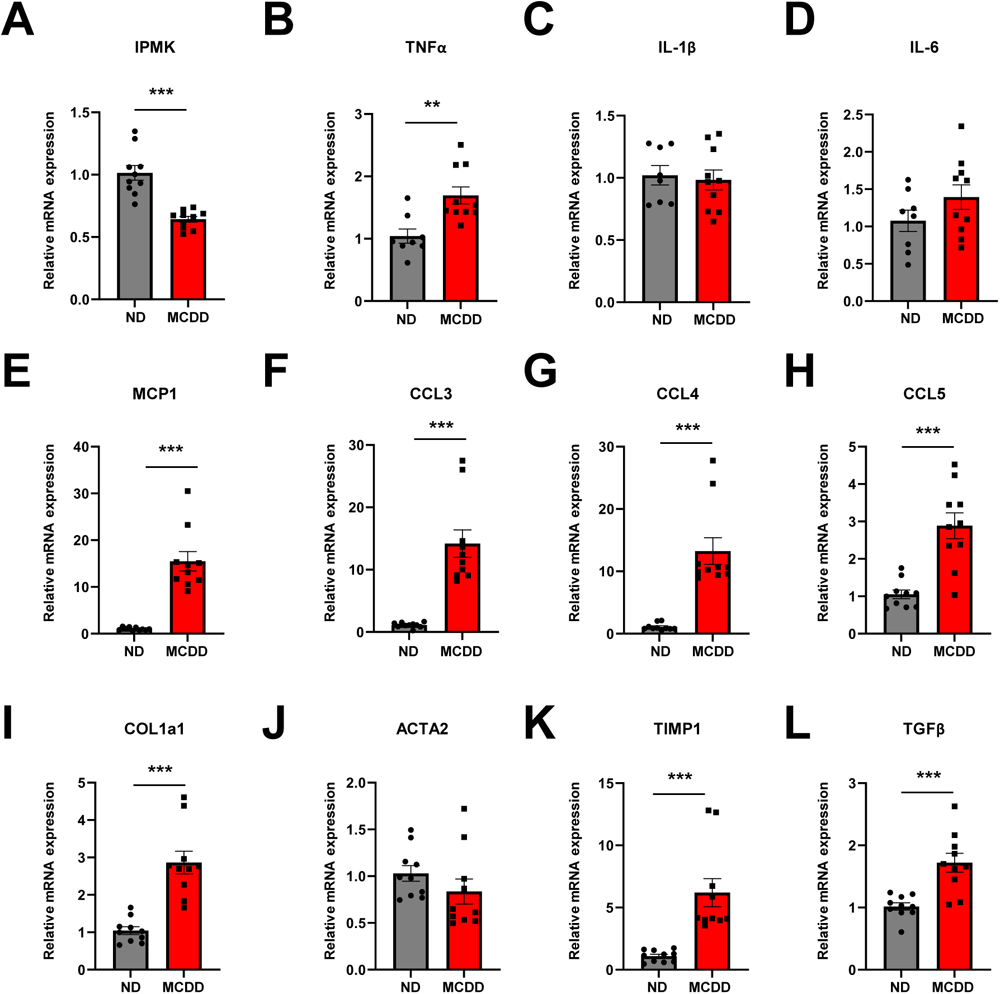
IPMK mRNA is decreased in the liver of MCDD mice. mRNA levels of (A) IPMK, (B-D) inflammatory cytokines, (E-H) chemokines, and (I-L) fibrosis-related genes in the liver from ND and MCDD-fed mice (n = 4-5 per group). Data are mean ± SEM; **p < 0.01, ***p < 0.001.

### Loss of hepatic IPMK exacerbates NASH progression

Since IPMK expression was significantly reduced in the liver of MCDD WT mice, we investigated whether IPMK is involved in NASH progression in a liver-specific IPMK knockout (LKO) mouse model. WT and LKO mice were fed MCDD for a short duration, 2 weeks, to induce steatohepatitis. Both the WT and LKO mice lost similar amounts of body weight (Fig. 3A); however, hepatic TG, serum ALT and AST levels were significantly increased in LKO compared to WT mice (Fig. 3B-D). Histological analysis revealed increased hepatic steatosis, inflammation, and fibrosis in the liver of LKO mice on MCDD (Fig. 3E). The expression levels of pro-inflammatory cytokines were not significantly different between WT and LKO mice (Fig. 4B-D), but pro-inflammatory chemokines were significantly increased in LKO liver of MCDD mice (Fig. 4E-H). These results suggest that hepatic IPMK deficiency promotes NASH progression.

**Figure 3.**
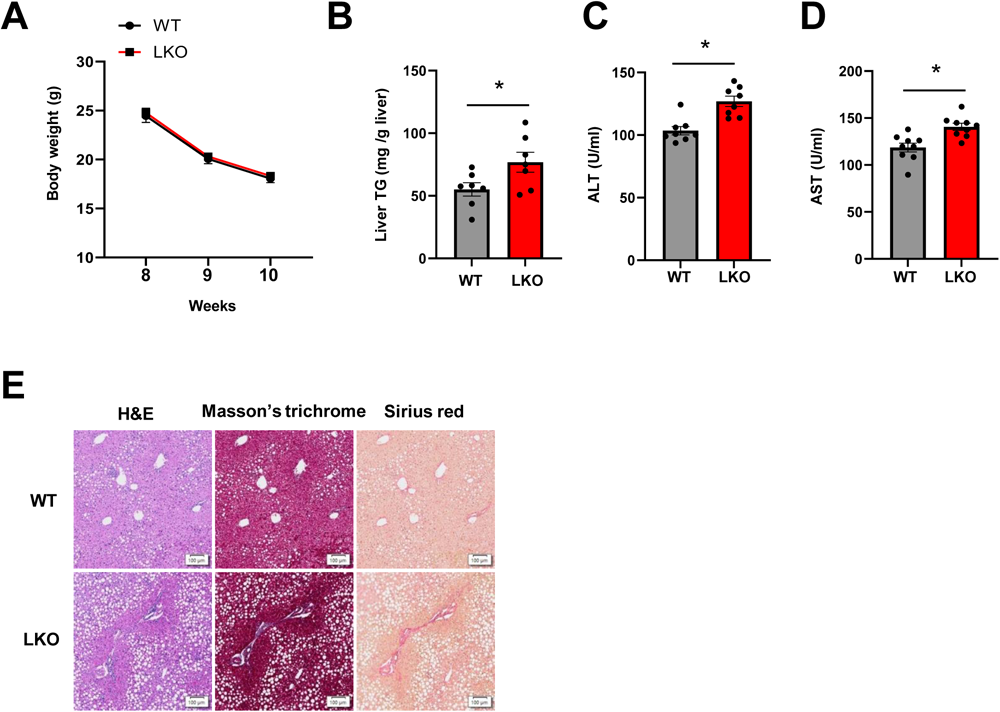
Loss of hepatic IPMK exacerbates NASH progression. (A) Body weight of WT (n = 5-8) and LKO (n = 5-9) mice fed MCDD was measured weekly from age 8 to 10 weeks. (B) Liver TG, serum (C) ALT and (D) AST were measured to evaluate the levels of liver injury. (E) Liver paraffin sections were stained with H&E, Masson’s trichrome, and Sirius red to determine the levels of steatosis, inflammation, and fibrosis (scale bar, 100 um). Data are mean ± SEM; *p < 0.05.

**Figure 4.**
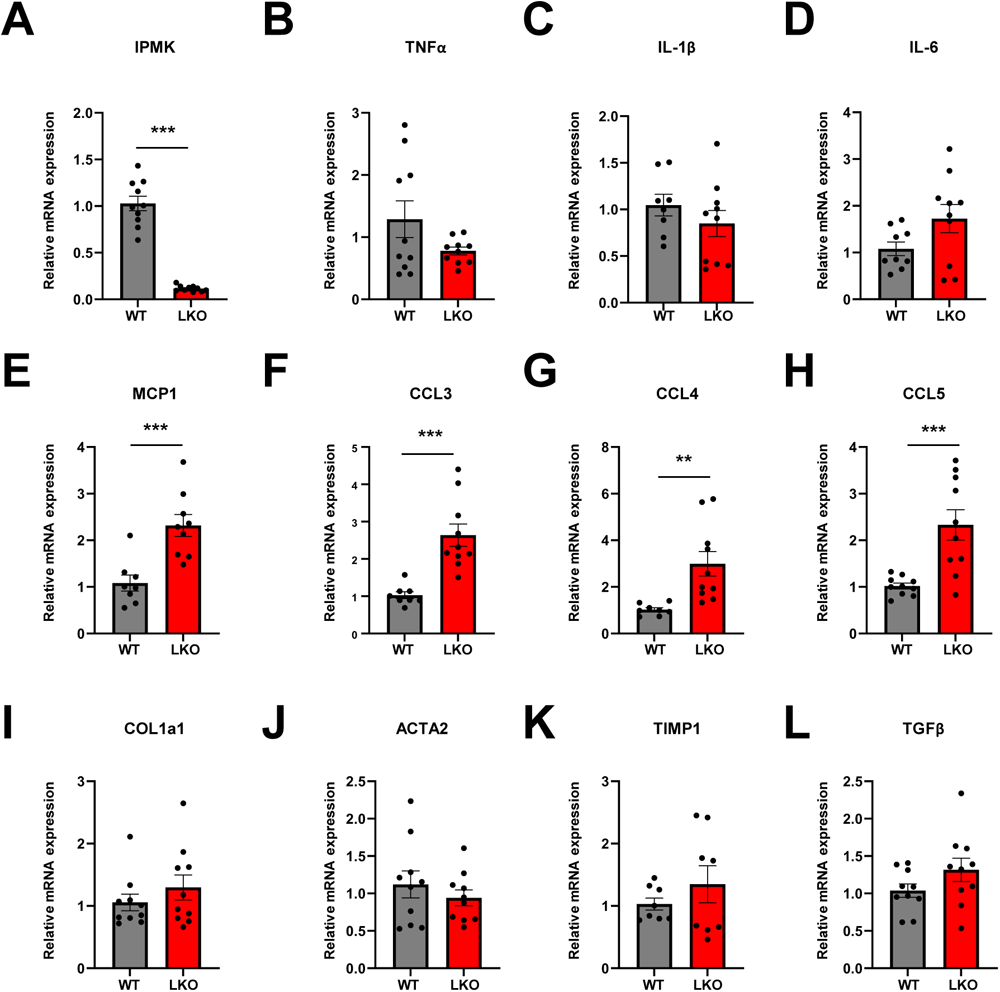
Loss of hepatic IPMK upregulates pro-inflammatory chemokine gene expression. mRNA levels of (A) IPMK, (B-D) inflammatory cytokines, (E-H) chemokines, and (I-L) fibrosis-related genes in the liver from MCDD WT and LKO mice (n = 4-5 per group). Data are mean ± SEM; **p < 0.01, ***p < 0.001.

### TRF prevents NASH progression and restores IPMK expression level

Many studies have shown that TRF has beneficial metabolic effects such as weight loss, improved insulin sensitivity, reduced hepatic steatosis and hyperlipidemia, and amelioration of inflammation (Chung et al., 2016; Marjot et al., 2023; Xie et al., 2022). We determined whether TRF could prevent the NASH phenotype induced by MCDD. To investigate the effects of TRF on NASH, 10-week-old mice were fed MCDD *ad libitum* (ALF) or were restricted to feeding from 7 pm to 8 am (TRF) for 2 weeks. Body weight was reduced in the TRF group compared to the ALF group (Fig. 5A), despite no significant change in food intake (Fig. 5F), and hepatic TG, serum ALT, and AST levels were reduced compared to ALF mice (Fig. 5B-D). Histological analysis showed decreased hepatic steatosis, inflammation, and fibrosis in the liver of TRF mice (Fig. 5E). The expression levels of pro-inflammatory cytokines (Fig. 6B-D), chemokines (Fig. 6E-H) and fibrosis related genes (Fig. 6I and K) were significantly reduced in the liver of TRF mice. IPMK protein levels were increased in the liver of TRF mice compared to ALF mice (Fig. 5G). These results suggest that TRF prevents NASH progression by maintaining hepatic IPMK levels.

**Figure 5.**
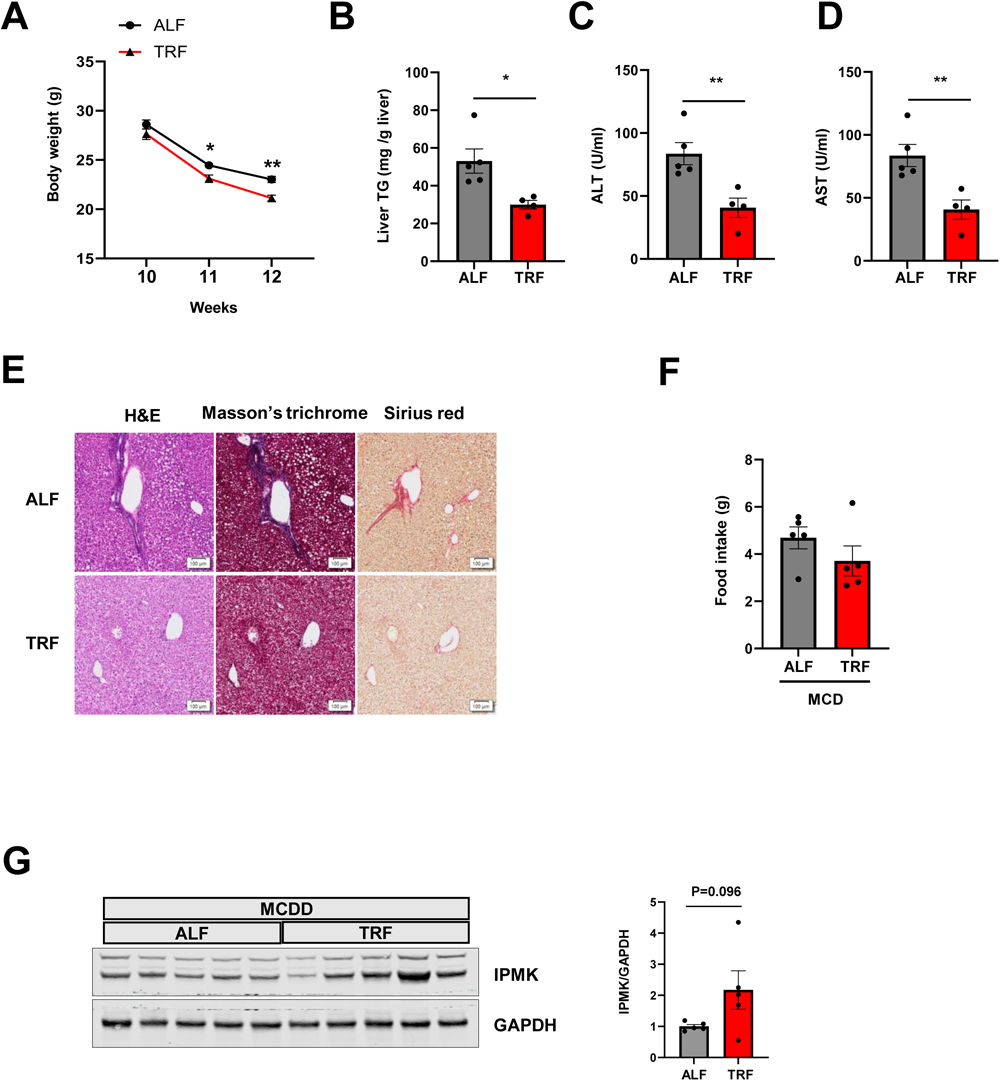
TRF prevents NASH progression. (A) Body weight of ad libitum fed (ALF) (n = 5) and time restricted fed (TRF) (n = 4-5) mice on MCDD were measured weekly from age 10 to 12 weeks. (B) Liver TG, serum (B) ALT and (D) AST were measured to evaluate the levels of liver injury. (E) Liver paraffin sections were stained with H&E, Masson’s trichrome, and Sirius red to determine the levels of steatosis, inflammation, and fibrosis (scale bar, 100 um). (F) Food intake (g/day) of ALF and TRF mice fed MCDD was measured. (G) Protein levels of IPMK in the liver from ALF and TRF mice were analyzed by immunoblotting (n = 5 per group). Levels of IPMK protein were quantified. Data are mean ± SEM; *p < 0.05, **p < 0.01.

**Figure 6.**
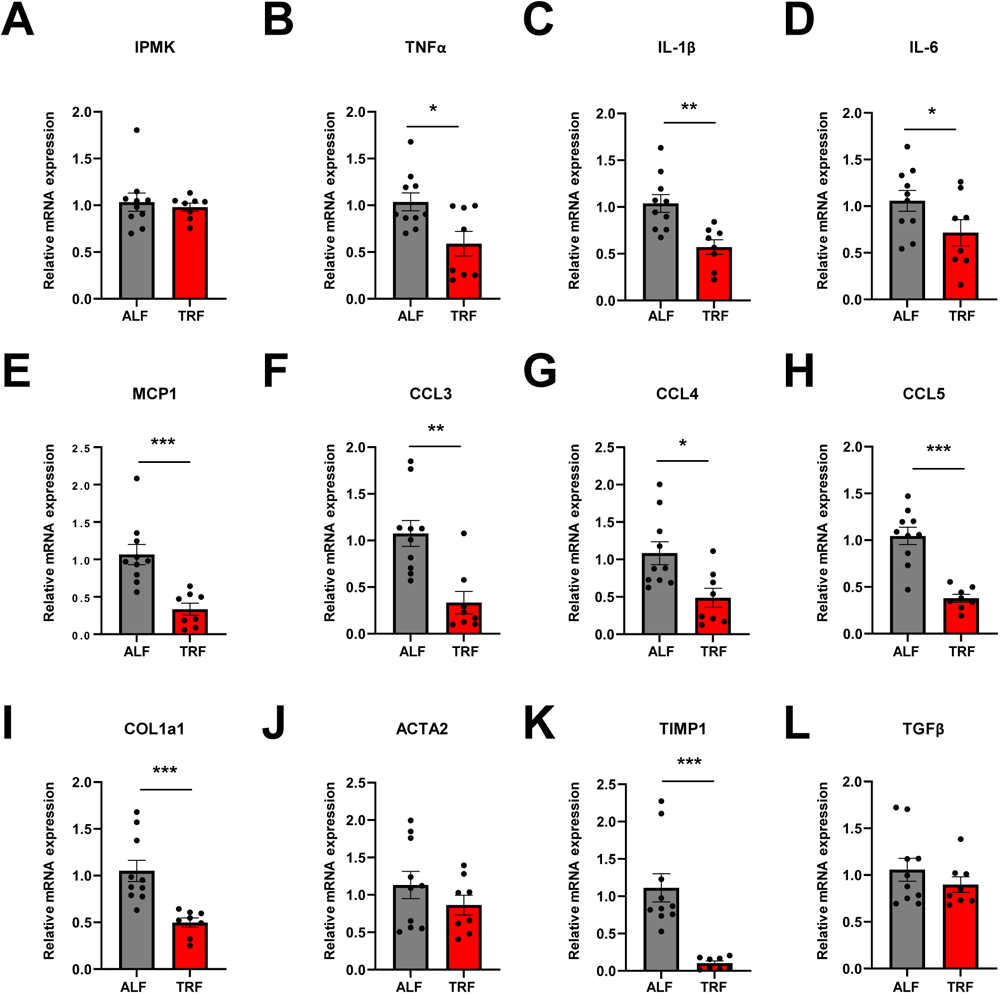
TRF prevents MCDD-induced pro-inflammatory gene expression. mRNA levels of (A) IPMK, (B-D) inflammatory cytokines, (E-H) chemokines, and (I-L) fibrosis-related genes were measured in the livers of both the ALF and TRF groups (n = 4-5 per group). Data are mean ± SEM; *p < 0.05, **p < 0.01, ***p < 0.001.

## Discussion

Nonalcoholic fatty liver disease (NAFLD) is a major public health challenge, affecting 20-30% of adults in the general population, and more than 70% of patients with obesity and diabetes (Wei et al., 2023). NAFLD may progress from simple steatosis to NASH, cirrhosis, liver failure, and hepatocellular carcinoma (Cholankeril et al., 2017). Dietary calorie restriction reduces weight and hepatic steatosis in people with NAFLD; however, the long-term adherence to calorie restriction is challenging. Time-restricted eating (TRE) is a popular intermittent fasting regimen involving a specific eating period within a 24-hour cycle. TRE has gained attention because it is easier to adhere to and has metabolic benefits (Lowe et al., 2020). Studies in rodents suggest that food timing rather than total calorie intake per se underlies the beneficial effects of TRE (Hatori et al., 2012). Although TRF reduces hepatic lipids and inflammation in mice, the underlying mechanisms are unknown (Chung et al., 2016; Hatori et al., 2012).

In this study, we demonstrated a mechanistic link between TRF, IPMK and NASH in a mouse model. Hepatic-specific deletion of IPMK exacerbates MCDD-induced steatohepatitis and fibrosis in mice. TRF prevents a decrease of IPMK expression by MCDD and attenuates MCDD-induced steatohepatitis. Many groups have examined the effects of TRF on metabolic parameters such as glucose regulation, insulin sensitivity, and body composition, in animals and humans (Chaix et al., 2014; Hatori et al., 2012; Sutton et al., 2018). The initial evidence supporting the metabolic benefits of TRF was gathered through experiments conducted on mice with diet-induced obesity, utilizing a high-fat diet (Hatori et al., 2012). Subsequent research has consistently used diets designed to induce obesity, including high-fat diets, high-fat plus high-sucrose, and high-fructose diets, in order to investigate the effects of TRF. However, no study to date has investigated the effects of TRF in the context of NASH induced by MCDD. Although MCDD has certain limitations, such as weight loss and absence of insulin resistance, it has been a commonly used dietary model for NASH in mice, which results in liver injury characterized by excessive fat accumulation, inflammation, and fibrosis (Ipsen et al., 2020; Rinella & Green, 2004; Rivera et al., 2007). In our study, we observed that TRF significantly reduced hepatic lipid accumulation and liver damage caused by the MCDD. Interestingly, despite the MCDD being a diet that promotes rapid weight loss, the TRF group actually showed greater weight reduction compared to the ad libitum-fed (ALF) group (Fig. 5A). This result is consistent with the weight reduction typically observed in TRF studies in mice (Hatori et al., 2012; Schroder et al., 2021).

Chemokines play an important role in maintaining tissue homeostasis and promoting inflammation in various organs, including the liver. Chemokines guide immune cells to the liver through specific chemokine receptors. Both liver-resident and infiltrating cells can release chemokines and cytokines in response to injury. Several chemokines are upregulated in liver and adipose tissues of patients with NAFLD (Braunersreuther et al., 2012). MCP1, produced by hepatocytes, stellate cells, and Kupffer cells during liver pathology, is a crucial chemokine that stimulates the activation of monocytes and macrophages (Sahin et al., 2010). Elevated levels of MCP1 have been linked to the progression of hepatic inflammation and fibrosis (Cao et al., 2021). CCL5 is another important chemokine in liver fibrogenesis (Berres et al., 2010). Patients with liver cirrhosis often exhibit increased levels of CCL5, along with related chemokines CCL3 and CCL4 (Berres et al., 2010; Sadeghi et al., 2015; Xu et al., 2021). In our MCDD NASH model, we showed that TRF significantly reduced inflammatory chemokines (Fig. 6E-H) as well as inflammatory cytokine gene expression (Fig. 6B-D) compared to *ad libitum* feeding group. Further investigation is necessary to evaluate how TRF specifically reduces chemokine gene expression induced by MCDD. It is possible that TRF modifies leukocyte responsiveness and improves inflammation (Y. Kim et al., 2022; Y. Lee et al., 2021).

Despite numerous studies on the beneficial effects of TRF, the molecular mechanisms have not been clearly elucidated. Many studies have focused on the role of circadian clock genes in metabolic architecture since feeding and fasting responses interact with the molecular circadian clock to affect diurnal rhythms (He et al., 2016; Koronowski & Sassone-Corsi, 2021; Takahashi, 2017; Turek et al., 2005). However, Chaix et al. reported that TRF still prevents obesity and metabolic syndrome in mice lacking a circadian clock, such as Cry1:Cry2, Bmal1, Rev-erva/b (Chaix et al., 2019). IPMK plays a crucial role in the metabolism of inositol polyphosphates (IPs) and functions as a versatile signaling factor intricately intertwined with essential biological pathways, including those related to AMPK, mTOR, and Akt ((Bang et al., 2012; S. Kim et al., 2011; Maag et al., 2011)). Additionally, IPMK has been associated with liver inflammation (Guha et al., 2019) and oxidative stress (Tyagi et al., 2023). Given that the development of NAFLD/NASH is linked to factors such as insulin resistance, lipid metabolism, oxidative stress, and inflammation, it is conceivable that IPMK may play a role in the development of NAFLD/NASH. Indeed, we found that IPMK expression was decreased by MCDD, and the loss of hepatic IPMK worsened steatohepatitis induced by MCDD. Interestingly, we found that TRF restored the MCDD-induced reduction in IPMK protein and pro-inflammatory chemokine gene expression, suggesting that IPMK may play a causal role in the progression of NAFLD/NASH.

Further studies are needed to understand how IPMK mediates changes in hepatic lipid metabolism, inflammation, and fibrosis in NASH, and how TRF modulates the levels of IPMK in the liver to prevent the development of NASH. Our study in MCDD mice presents a model for elucidating the pathophysiology and potential treatment of NAFLD/NASH.

## Materials and Methods

### Animals

Experiments were performed in accordance with the Institutional Animal Care and Use Committee guidelines with its approvals. The LKO mice were generated by crossing Albumin-CreTg/+ mice (Jackson Laboratories) with mice homozygous for a “floxed” exon 6 of IPMK (IPMK *fl//fl*). The control mice for this study were IPMK *fl/fl* (WT) mice. Mice were housed under standard conditions in a temperature-and humidity-controlled facility with a light–dark cycle of 12 h (lights on at 07:00) and fed ad libitum at the Johns Hopkins University (Baltimore, MD). WT and LKO mice aged 8 weeks were fed a ND (A02082003BY, Research Diets, New Brunswick, NJ, USA) or MCDD (A02082002BR, Research Diets, New Brunswick, NJ, USA) for 2 weeks.

### Liver Triglyceride measurement

Liver triglyceride was extracted using the Folch extraction method (Folch et al., 1957), and the triglyceride levels were measured using StanbioTriglyceride LiquiColor kit (Stanbio Laboratory, Boerne, TX, USA) according to manufacturer’s protocol.

### ALT and AST measurement

Mice were euthanized and blood obtained from cardiac puncture and centrifuged (2,000g, 10 min) at 4°C. The serum layer was collected and stored at −80°C. Serum ALT and AST were measured using the ALT/SGPT Liqui-UV® test and AST/SGOT Liqui-UV® test (Stanbio Laboratory, Boerne, TX, USA) according to the manufacturer’s protocol.

### Liver Histology

Liver tissues fixed with PBS containing 4% paraformaldehyde were embedded in paraffin, and 5-μm sections were stained with hematoxylin and eosin solution, Masson’s trichrome solution or Sirius Red solution, according to standard procedures.

### Immunoblotting

Liver tissues were lysed in RIPA buffer containing inhibitors and heated at 95°C for 5 min prior to electrophoresis. Proteins were transferred to a 0.2-mm nitrocellulose membrane, blocked with 5% nonfat dry milk, and incubated with primary antibodies at 4°C overnight. Immunoblotting was conducted with the following antibodies: IPMK from Novus Biologicals, GAPDH from Cell Signaling Technology. Blots were imaged and quantitated using an Odyssey Near-Infrared Scanner (Li-Cor Biosciences, Lincoln, NE, USA).

### Quantitative real-time PCR

Total RNA was isolated from WT or LKO mouse liver tissue or primary hepatocytes using TRIZol reagent (Thermo Fisher Scientific, Waltham, MA, USA), and chloroform (MilliporeSigma, Burlington, MA, USA), precipitated in 2-propanol (MilliporeSigma, Burlington, MA, USA), washed in ethanol (MilliporeSigma, Burlington, MA, USA), and quantified using Epoch Microplate Spectrophotometer (Agilent Technologies, Santa Clara, CA, USA). mRNA was reverse transcribed into cDNA using ProtoScript® II First Strand cDNA Synthesis Kit (BioLabs, Ipswich, MA, USA). Ribosomal Protein L32 (RPL32) mRNA was used as the invariant endogenous control and melting curve analysis was run to verify specificity of each amplicon. The relative amounts of the RNAs were calculated using the comparative threshold cycle method.

### Statistical Analysis

Statistical analyses were performed using GraphPadPrism 10 (GraphPad, San Diego, CA, USA). Data are presented as mean ± SEM. Comparisons between two groups were done using Student’s t test. The threshold for statistical significance was set at P < 0.05.

## Author contributions

SFK and IJ designed and directed the project. IJ performed the experiment. IJ analyzed data and wrote the first draft of the manuscript. RSA and SFK reviewed and edited the manuscript. All authors approved the paper for submission.

## Acknowledgments

This work was supported by grants from American Heart Association Strategically Funded Research Network (SFRN) Obesity Center (RSA and SFK, 17SFRN33610014 and 20SFRN35210662), National Institutes of Health/National Institute of Diabetes and Digestive and Kidney Diseases (NIH/NIDDK) (SFK and RSA, DK135751), and American Heart Association SFRN fellowship (IJ, 17SFRN33560006).

